# Shear flow-driven actin re-organization induces ICAM-1 nanoclustering on endothelial cells that impact T-cell migration

**DOI:** 10.1101/2020.06.29.177816

**Authors:** Izabela K. Piechocka, Sarah Keary, Alberto Sosa-Costa, Lukas Lau, Nitin Mohan, Jelena Stanisavljevic, Kyra J. E. Borgman, Melike Lakadamyali, Carlo Manzo, Maria F. Garcia-Parajo

## Abstract

The leukocyte specific β_2_-integrin LFA-1, and its ligand ICAM-1 expressed on endothelial cells (ECs), are involved in the arrest, adhesion and transendothelial migration of leukocytes. Although the role of mechanical forces on LFA-1 activation is well established, the impact of forces on its major ligand ICAM-1, has received less attention. Using a parallel-plate flow chamber combined with confocal and super-resolution microscopy, we show that prolonged shear-flow induces a global translocation of ICAM-1 on ECs upstream of flow direction. Interestingly, shear-forces promoted ICAM-1 nanoclustering prior to LFA-1 engagement. This spatial nanoscale organization was driven by actin cytoskeleton re-arrangements induced by shear-force. We further assessed the impact of prolonged shear-stress EC stimulation on T cell migration. T cells adhered to mechanically pre-stimulated ECs developed a more pro-migratory phenotype, migrated faster and exhibited shorter EC interactions than when adhered to non-mechanically stimulated ECs. Together, our results indicate that shear-forces increase the number of ICAM-1/LFA-1 bonds due to ICAM-1 nanoclustering, strengthening adhesion and thereby reducing actin retrograde flow of T-cells, leading to their increased migration speed. Our data also underscores the importance of mechanical forces regulating the spatial organization of cell membrane receptors and their contribution to adhesion regulation, regardless of integrin activation.

**Summary statement:** We show that shear forces promote ICAM-1 spatial re-arrangement and actin-dependent nanoclustering on ECs prior to integrin engagement. This mechanism might be important for firm leukocyte adhesion and migration during the immune response.

## INTRODUCTION

Endothelial cells (ECs) form a natural barrier for regulating leukocyte migration within blood vessels. Leukocyte extravasation from the bloodstream to sites of inflammation or peripheral lymphoid organs involves a cascade of steps of leukocyte interactions with the activated endothelium. This multistep process includes initially low-affinity adhesive interactions (capture and rolling), firm adhesion, cell spreading and crawling, and finally, leukocyte diapedesis through the endothelial barrier to the sites of inflammation (Ley et al., 2007; Nourshargh and Alon, 2014). These processes are mainly mediated by interactions between the leukocyte specific α_L_β_2_ integrin LFA-1, and its major ligand, the Intracellular Adhesion molecule (ICAM-1), which is highly expressed on activated ECs (Imhof and Aurrand-Lions, 2004; Makgoba et al., 1988; Marlin and Springer, 1987). The various degrees of adhesion during this cascade of events requires thus a tight regulation of LFA-1 activation and its binding strength to ICAM-1.

It is well-known that LFA-1 activation is mediated by outside-in or inside-out triggering events (Abram and Lowell, 2009; Hogg et al., 2011; Kinashi, 2005; Luo et al., 2007). Avidity and lateral mobility also contribute to integrin-mediated adhesion (Bakker et al., 2012; van Kooyk and Figdor, 2000). Moreover, tensile mechanical forces exerted by fixed ICAM-1 ligands have been recently proposed to stabilize LFA-1 in an active conformation (Hogg et al., 2011; Nordenfelt et al., 2016; Schürpf and Springer, 2011). On circulating leukocytes, lateral shear-forces provided by flow conditions lead to full activation of LFA-1 through the opposing forces exerted by immobilized ICAM-1 and the actin cytoskeleton (Chen et al., 2010; Hogg et al., 2011). Such force-induced changes in integrin conformation expose further ligand-binding sites reinforcing the initially created integrin-ligand bonds and leading to increased leukocyte binding to the endothelium (Constantin et al., 2000; Li et al., 2018; Shamri et al., 2005). However, most of these experiments have been performed on substrates containing immobilized ICAM-1 since soluble ligands are not able to activate integrins as they do not provide the additional counter force required to stabilize integrins in their active form (Chen et al., 2010; Hogg et al., 2011; Li et al., 2018). Intriguingly, mobility measurements of unengaged ICAM-1 showed that a large percentage of the ligand freely diffuses on the EC membrane (van Buul et al., 2010). These findings are difficult to conciliate with the requirement of ICAM-1 immobilization and tensile forces to fully activate LFA-1.

Due to their localization on the EC surface, ICAM-1 molecules are continuously exposed *in-vivo* to fluid shear stresses generated by blood flow. Together with their counter integrin receptors, these ligand-integrin complexes therefore not only have to withstand traction forces exerted by crawling leukocytes but also have to resist forces generated by flowing blood. It has been previously shown that shear forces have an impact on ECs by inducing cell shape and actin cytoskeleton organization (Galbraith et al., 1998; Wojciak-Stothard and Ridley, 2003), focal adhesion formation (Davies et al., 1994), changes in membrane fluidity (Butler et al., 2001; Yamamoto and Ando, 2013) and by modulating gene regulation (Nakajima and Mochizuki, 2017). Moreover, earlier reports showed that shear-forces upregulate the expression levels of ICAM-1 (Nagel et al., 1994; Tsuboi et al., 1995). More recent work indicates that ICAM-1 is a force-sensor that can initiate mechanosensitive signaling events that include actin-cytoskeleton re-organization, myosin-based contractile forces and EC stiffness (Schaefer and Hordijk, 2015).

Structurally, ICAM-1 contains five extracellular glycosylated Ig-like domains, a transmembrane domain and a short cytoplasmic tail (Staunton et al., 1990). It has been also shown that ICAM-1 structurally exist as a dimer (Yang et al., 2004). Although the cytoplasmic tails contain no actin binding motifs, clustering of ICAM-1 (induced by anti-ICAM-1 Ab-coated beads or leukocyte binding) is capable of triggering the recruitment of cytoplasmic actin-binding proteins that anchor ICAM-1 to the actin cytoskeleton and increase its immobilization (Schaefer et al., 2014; Schaefer and Hordijk, 2015; van Buul et al., 2010). Similarly, actin-regulated ICAM-1 clustering and restricted mobility have been observed on dendritic cells upon maturation (Comrie et al., 2015). Yet, the lateral mobility and spatial distribution of ICAM-1 in the *presence of shear-force and prior to leukocyte engagement* has not been studied in detail. It is also less known how prolonged pre-exposure of ECs to fluid shear-stress affects leukocyte migratory behavior. To address these questions, we pre-exposed inflammatory challenged ECs to continuous shear flow prior to T-cell engagement and visualized over time the spatial distribution of ICAM-1 and the actin cytoskeleton on ECs. We show that ICAM-1 molecules are part of the shear-sensitive EC machinery that rearranges in response to continuous flow, forming nanoclusters on the EC surface that are highly localized upstream of flow. This particular spatial organization is driven by actin-cytoskeleton re-arrangements induced by shear-forces. ICAM-1 nanoclustering might thus contribute to strengthening interactions with integrins expressed on T-cells and adhered to ECs, allowing T-cells to exert larger traction forces needed for migration over the endothelium.

## RESULTS

### Prolonged shear stress induces global translocation of ICAM-1 upstream of flow regardless of inflammatory EC activation

To decouple the effect of mechanical forces from biochemical stimulation of ECs, we first investigated the effect of shear stress on ICAM-1 expression and distribution on ECs i.e., without biochemical stimulation, at the single cell level. We exposed near-confluent resting ECs to a continuous shear flow of 8 dyn cm^-2^, fixed the cells at different time points after prolonged shear flow stimulation, labeled ICAM-1 and visualized its overall distribution on individual cells by confocal microscopy (Fig. S1). In static conditions, ICAM-1 was barely detectable and uniformly distributed through the entire cell (Fig. S1). Application of shear stress resulted in a two-fold increase of the ICAM-1 signal over the course of six hours (Fig. S1). These results are consistent with earlier flow cytometry studies showing that shear-forces are capable on their own to upregulate ICAM-1 levels on ECs (Nagel et al., 1994; Tsuboi et al., 1995).

To test the effect of prolonged shear flow exposure on the expression levels and spatial distribution of ICAM-1 in the presence of inflammatory conditions, we treated ECs with TNFα, an inflammatory cytokine known to upregulate ICAM-1 expression. As expected, TNFα treatment resulted in translocation of ICAM-1 to the cell membrane, mostly to the apical side (Fig. 1A and z-projections) and its *de novo* expression, with a six-fold increase in the total ICAM-1 intensity as compared to resting ECs (Fig. 1B). Moreover, application of shear flow for up to six hours induced a modest but significant increase in ICAM-1 expression (Fig. 1C) indicating an additive effect of TNFα and prolonged shear stress on ICAM-1 expression.

**Fig. 1.**
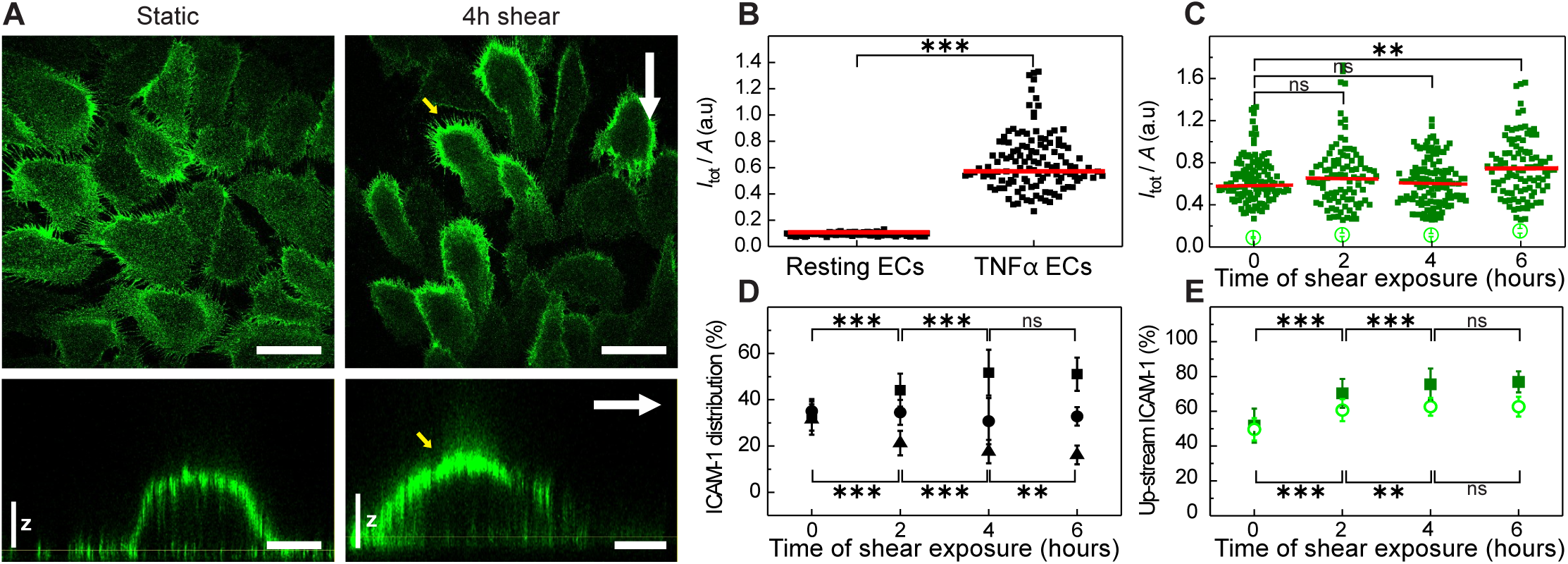
Shear flow induces global translocation of ICAM-1 upstream of flow regardless of inflammatory EC activation. (A) Single plane confocal microscopy images taken at the basal cell membrane (*top rows*) and 3*D* orthogonal views *(bottom rows*) of ICAM-1 expressed on TNFα-stimulated ECs in the absence (*lef*t) and presence of 4 hours continuous shear flow (*right*). White arrows indicate the direction of shear flow. Yellow arrows point at the upstream accumulation of ICAM-1 upon shear-force application. Scale bars, 50 μm in *top images* and 15 μm in *bottom images*. The vertical scale *z* is 3µm. (B) Fluorescence intensity of the ICAM-1 signal integrated over the entire cell and normalized to the cell area, for resting and for TNFα-stimulated ECs in static conditions, i.e., without flow application. Red lines indicate the mean fluorescence value. (C) Fluorescence intensity of the ICAM-1 signal integrated over the entire cell and normalized to the cell area, as a function of shear-flow time exposure. Time = 0 hours correspond to static conditions. Open symbols show the average ICAM-1 signal obtained over multiple resting ECs, while closed symbols correspond to individual TNFα-stimulated ECs. Red lines indicate the mean fluorescence value. (D) Percentage of up-(squares) middle-(circles) and down-stream (triangles) ICAM-1 fluorescence signal taken from individual TNFα-stimulated ECs, as a function of the shear-force application. The intensity was estimated from 33% of the desired cell area and divided by the total intensity calculated over the entire cell. (E) Percentage of upstream ICAM-1 fluorescence signal taken from individual ECs under resting (*open symbols*) and TNFα-stimulated (*closed symbols*) conditions, as a function of shear-force application. The signal was estimated from 50% of the desired cell area and divided by the total intensity calculated over the entire cell. All data are taken for n=100-180 ECs from 2 separate experiments per condition. Results are mean ± s.d. *** *p* < 0.001; ** *p* = 0.01; ns., not significant.

Interestingly, shear-flow induced a strong gradient on ICAM-1 distribution on the apical cell membrane, with highest ICAM-1 levels upstream of flow (Fig. 1A-*yellow arrows*). This upstream accumulation was dependent on flow exposure time, being already significant at two hours, and reaching a plateau after four hours of shear stimulation (Fig. 1D,E). A similar trend was also observed on resting ECs (i.e., without TNFα stimulation) (Fig. 1E-*open symbols*) indicating that ICAM-1 re-location upstream of flow was solely due to shear-force stimulation. Together, these data show a synergistic effect of biochemical and mechanical stimulation on the expression and spatial distribution of ICAM-1 on ECs. Whereas TNFα up-regulates ICAM-1 expression and induces translocation to the apical membrane, shear stress leads to a pronounced gradient in ICAM-1 apical re-distribution with highest levels located upstream of flow.

### Prolonged shear stress induces differential actin-cytoskeleton re-arrangements upstream and downstream of flow

It has been extensively documented that application of shear stress induces elongation and formation of actin fibers aligned to the flow direction (Galbraith et al., 1998). Confocal images of the actin cytoskeleton on ECs in the absence, and after 4 hours of shear flow stimulation confirmed cell elongation and actin cytoskeleton re-arrangements (Fig. 2A, Fig. S2). Moreover, most of the actin fibers aligned along the flow direction, but were preferentially located downstream of the flow (Fig. 2A, enlarged images). In contrast, a finer actin mesh, difficult to resolve by diffraction-limited confocal microscopy, appeared upstream of the flow direction (Fig. 2A). To gain more information on these finer actin structures, we performed super-resolution imaging by means of stochastic optical reconstruction microscopy (STORM) (Fig. 2B). With an effective increased resolution of ∼25nm, we clearly resolved the formation of actin-patch structures located preferentially upstream of the flow direction after four-hours of shear-flow exposure (Fig. 2B, *small white arrows*). The area occupied by these actin-patches were between 1-5 μm^2^, which is at least two-fold larger than any small actin puncta observed in static ECs (Fig. 2B and Fig. S3). Interestingly, the location of these actin patches coincided with regions of high ICAM-1 accumulation, prompting us to investigate the potential spatial relationship between them by confocal and super-resolution STORM.

**Fig. 2.**
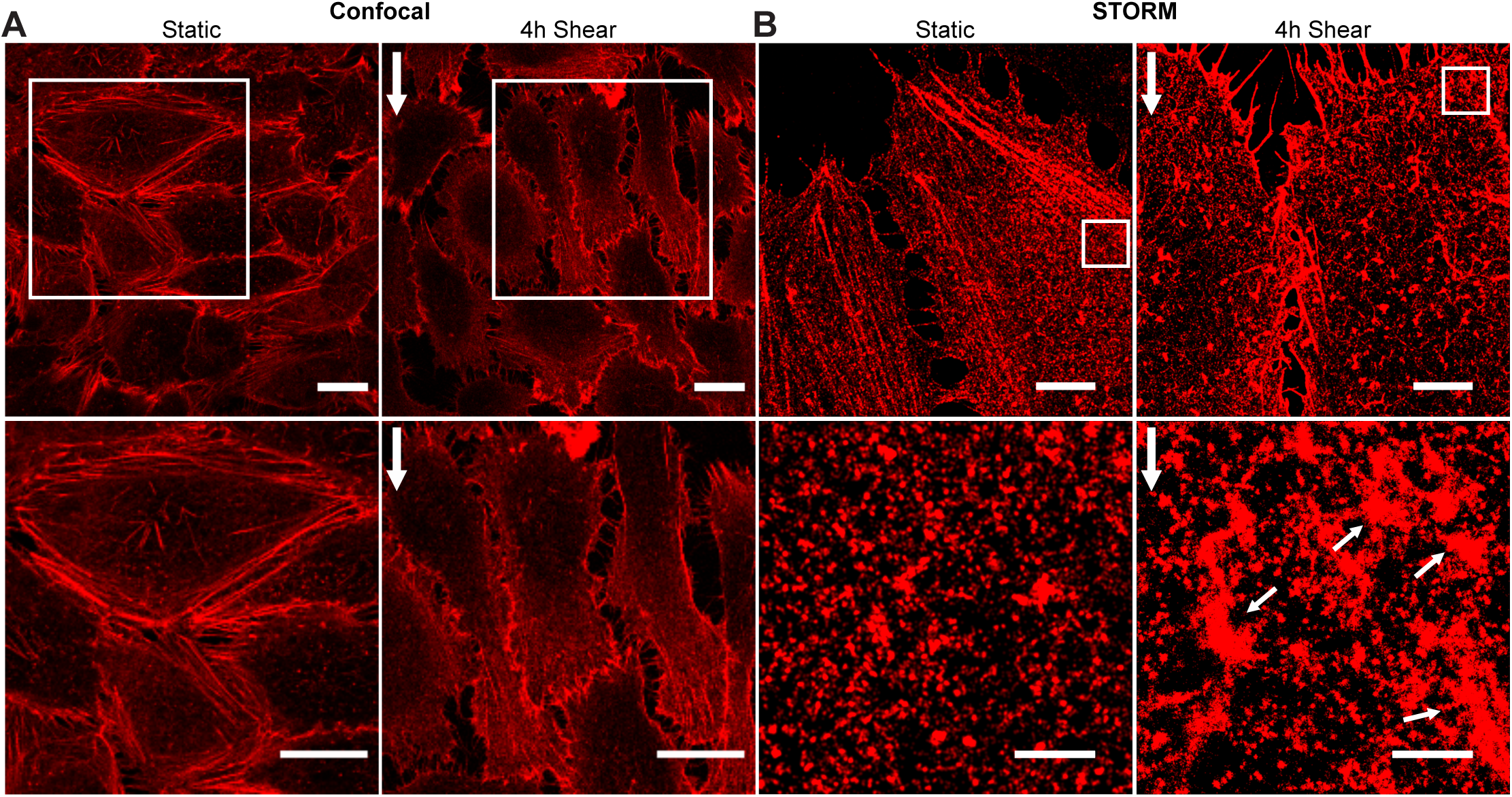
Prolonged shear stress induces differential actin-cytoskeleton re-arrangements upstream and downstream of flow. (A) Representative *Z*-projection confocal microscopy images (*top*) and magnified images (*bottom*) of the actin cytoskeleton on TNFα-stimulated ECs in the absence (*lef*t) and presence of 4 hours continuous shear flow (*right*). Vertical white arrows on the right indicate the flow direction. Scale bars, 20 μm. (B) Representative STORM images (*top*) and corresponding magnified images (*bottom*) of the actin cytoskeleton in static TNFα-treated ECs (*left*) and after 4 hours of shear-flow exposure (*right*). Vertical white arrows indicate the direction of the flow. Small white arrows point to actin patch-like structures. Scale bars, 5 μm (*full images*) and 1 μm (*zoom regions*).

### Patch-like actin structures localize with ICAM-1 rich regions upstream of flow

We first used dual color confocal microscopy to investigate the spatial correlation of actin and ICAM-1 in cell regions upstream of flow direction. We identified highly dense regions of actin patches decorated with ICAM-1 after four hours of shear-flow stimulation, as compared to the more discrete puncta detected in static conditions (Fig. 3A). The patch-like actin structures were not observed in downstream cell regions (Fig. S4). Image quantification showed more than a two-fold increase in the surface area occupied by newly formed actin-like patches (Fig. 3B) and the intensity of ICAM-1 concomitantly increased in these regions as compared to static conditions (Fig. 3C). These changes were accompanied by an increase in the overall colocalization between both structures (Pearson coefficient 0.47 *vs*. 0.34, for shear-flow and static conditions, respectively, Fig. S5).

**Fig. 3.**
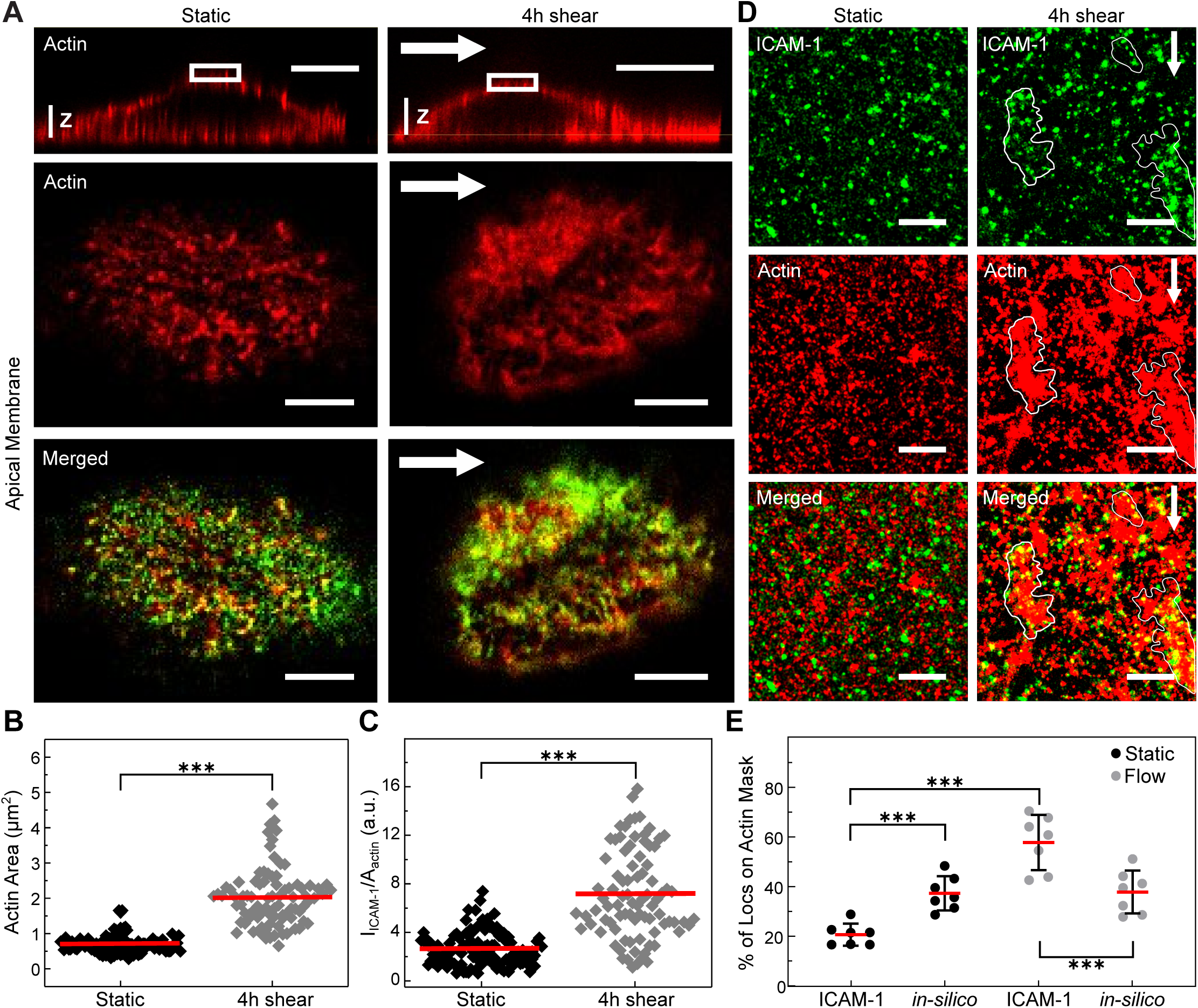
Shear flow promotes the formation of patch-like actin structures that localize with ICAM-1 rich regions upstream of flow. (A) 3*D* orthogonal views (*top row*) and magnified single plane confocal microscopy images (*middle* and *bottom rows*) of the cortical actin cytoskeleton (*red*) and ICAM-1 (*green*) localized at the apical cell membrane in static (*left*) and 4 hours shear-flow TNFα-stimulated ECs (*right*). White arrows indicate the direction of flow. Scale bars, 20 μm (*top images*) and 5 μm (*bottom images*), respectively. The vertical scale *z* is 3 µm. (B) Area occupied by puncta-like or patch-like actin structures, calculated from manually outlining actin regions. (C) ICAM-1 fluorescent intensity normalized to the corresponding actin area. Red lines indicate the mean fluorescence value (n=30 ECs from 2 separate experiments per condition). (D) Dual color STORM images on TNFα-stimulated ECs of ICAM-1 (*top*), actin (*middle*), and merged images (*bottom*) located upstream of the cells, in the absence (*left*) and after 4 hours shear-flow (*right*). White arrows show the direction of applied shear flow. White contours highlight regions where ICAM-1 co-localizes with actin patch-like structures. Scale bar, 1 μm. (E) Percentage of ICAM-1 single-molecule localizations residing inside actin, for static and 4 hours stimulated ECs. Symbols correspond to the calculated percentage per individual region of interest analyzed. *In-silico* data correspond to similar analysis but performed by randomly distributing on the actin mask the same number of ICAM-1 localizations as obtained from the experimental data. *** *p* < 0.001.

To gain more insight into the nanoscale re-arrangement of both actin and ICAM-1, we applied dual-color STORM. As this technique works best under total internal reflection illumination, we restricted our imaging to cortical regions located just above the basal cell membrane and upstream of flow. Concomitant to the reorganization of the actin cytoskeleton, we observed numerous discrete ICAM-1 spots, indicative of ICAM-1 nanoclustering (Fig. 3D). Moreover, a large majority of the ICAM-1 spots appear to reside inside or in close proximity to the shear-induced actin patches (Fig. 3D-*white outlines*). To quantify these results, we generated masks of the STORM actin signal, super-imposed to them the single molecule localizations events obtained on the ICAM-1 channel, and calculated the percentage of ICAM-1 localizations inside the actin mask (Fig. S6). The results confirm a significant increase of ICAM-1 co-localization with actin patches after application of flow, as compared to non-stimulated cells (Fig. 3E). The results were further validated by *in-silico* simulations of random ICAM-1 localizations with respect to the experimentally generated actin mask (Fig. 3E). Overall, these data reveal a clear shear-flow induced actin reorganization upstream of flow direction that is accompanied by selective ICAM-1 recruitment to these dense actin regions.

### Shear-flow induces the formation of actin-dependent ICAM-1 nanoclusters on TNFα stimulated ECs

Our STORM data showed the presence of discrete ICAM-1 spots on the cell membrane suggesting the formation of ICAM-1 nanoclusters. To further investigate and quantitate potential changes in ICAM-1 nanoscale organization as a result of shear-flow we used super-resolution stimulated emission depletion (STED) microscopy, which is more amenable to apical membrane studies. With a spatial resolution of ∼ 90 nm, STED resolved individual ICAM-1 spots packed at different densities, both in static conditions and after four hours of shear-flow stimulation (Fig. 4A-C). More sparse spots were observed in the absence of shear-flow (Fig. 4A) or in regions downstream of flow (Fig. 4C), whereas ICAM-1 spot density was much higher in regions of the membrane upstream of flow (Fig. 4B).

**Fig. 4.**
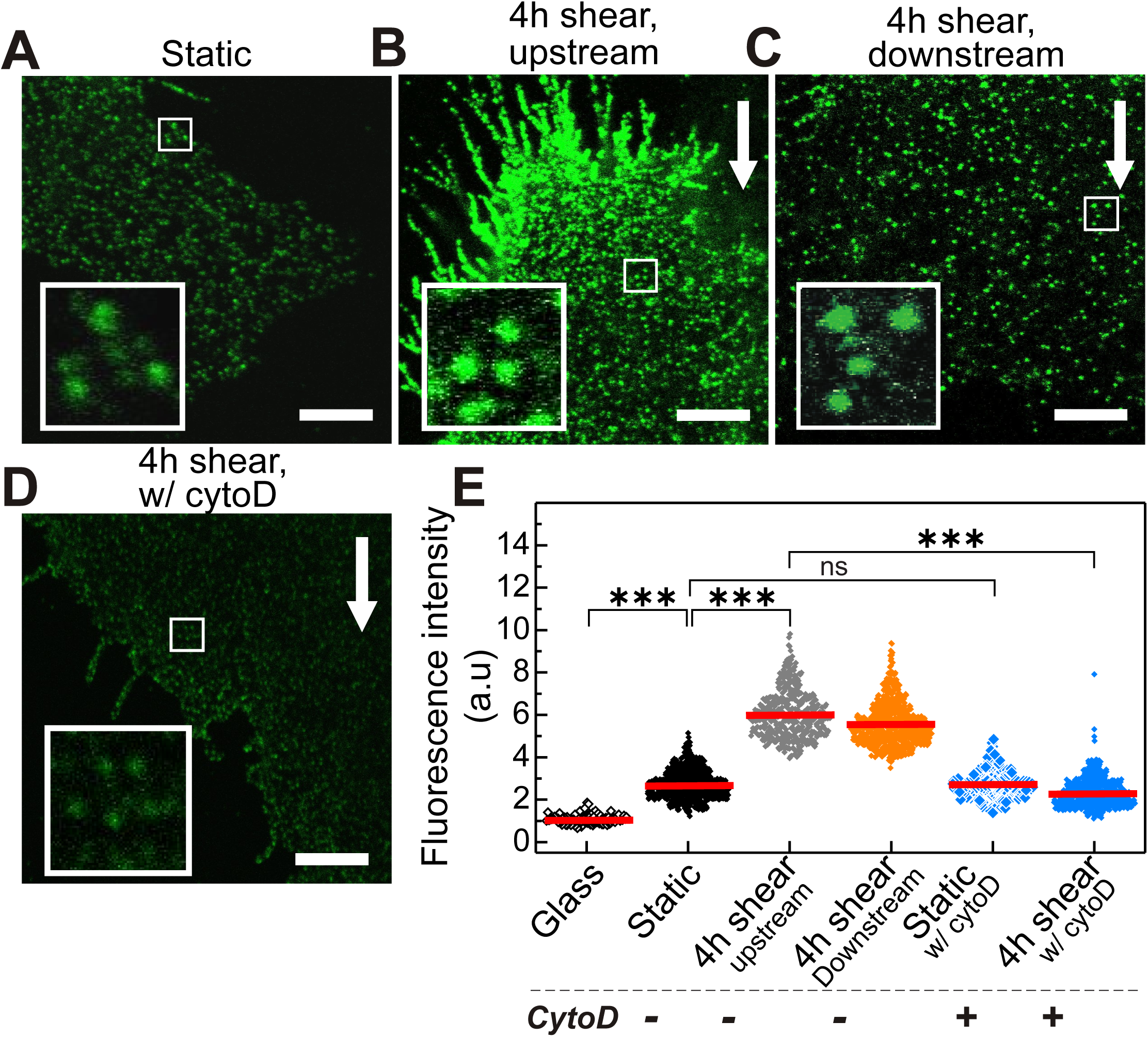
Shear flow promotes actin-dependent ICAM-1 nanoclustering on TNFα-treated ECs. Representative STED images of ICAM-1 taken in (A) static ECs, (B) after 4 hours of shear-flow stimulation, upstream of flow, (C) after 4 hours of shear-flow stimulation, downstream of flow and (D) after 4 hours shear-flow stimulated ECs and 20 minutes of CytoD treatment, upstream of flow. White arrows indicate the flow direction. Scale bars, 4 μm. (*Insets*) Magnified views of ICAM-1 fluorescent spots (1.5 x 1.5 μm regions). (E) Normalized ICAM-1 intensity per spot for different conditions, as extracted from the analysis of the STED images. The intensity of ICAM-1 has been normalized to the mean intensity of individual Abs non-specifically attached to the glass substrate. Red lines indicate the mean intensity values (n=30 cells from 2 independent experiments per condition). *** *p* < 0.001; ns., not significant.

To quantify the extent of ICAM-1 nanoclustering, we measured the fluorescence intensity of individual ICAM-1 spots over multiple STED images and compared it to that of sparsely scattered single fluorescence antibodies on the glass substrate. As intensity is proportional to the number of molecules, a large number of occurrences at high intensity values on the cell surface compared to single antibodies on glass indicates the occurrence of multiple molecules in each spot, i.e., nanoclustering (van Zanten et al., 2010). Under static conditions, the spots intensity showed on average a 2.7-fold increase in intensity as compared to individual fluorescence spots sparsely located on the glass coverslip (Fig. 4E), in agreement with the dimeric structure of ICAM-1 (Yang et al., 2004). Remarkably, application of shear flow resulted in a six-fold increase in spot intensity, demonstrating the formation of ICAM-1 nanoclusters on the apical EC membrane, both upstream and downstream of flow (Fig. 4E).

To enquire whether the observed ICAM-1 nanoclustering depends on the actin cytoskeleton, we used a mild concentration of cytochalasin D (CytoD) to disrupt the actin meshwork after shear force stimulation, and then imaged ICAM-1 distribution by STED. CytoD visibly perturbed the EC actin meshwork (Fig. S7) and caused marked changes in ICAM-1 nanoscale distribution (Fig. 4D). Indeed, perturbation of the actin cytoskeleton completely abrogated ICAM-1 nanoclustering with fluorescence spots having intensities comparable to those of static conditions (Fig. 4E). Together, these results demonstrate that shear-forces induce ICAM-1 nanoclustering prior to leukocyte engagement and underscore the crucial role of the actin cytoskeleton in forming and maintaining these nanoclusters. Of note, CytoD treatment did not affect ICAM-1 dimer distribution on static TNFα treated ECs (Fig. 4E), a result that is fully consistent with the notion that ICAM-1 is constitutively expressed as a dimer on the EC surface (Yang et al., 2004).

### Changes in ICAM-1 spatial organization as a result of shear-flow correlate with increased leukocyte migration across ECs

Upon inflammation, leukocytes migrate across the endothelium prior to extravasation in a process that is primarily mediated by interactions between ICAM-1 on the endothelium and its counterpart integrin receptor LFA-1 on the leukocyte membrane (Hogg et al., 2011; Nourshargh and Alon, 2014). To study the potential relevance of shear-flow induced ICAM-1 gradient distribution and its nanoclustering on the arrest and migration of leukocytes, we performed two sets of parallel experiments (Fig. 5A): one in which TNFα-treated ECs were first pre-stimulated for four hours by shear-flow, and the second set of experiments in which the cells were not mechanically pre-stimulated (i.e., static). For both experimental conditions, we then flowed Jurkat T-cells into the chamber, allowed them to settle on the ECs for three minutes, resumed the flow at 1 dyn cm^-2^ (post-flow) and recorded T-cell movement for 20 minutes (Fig. 5A).

**Fig. 5.**
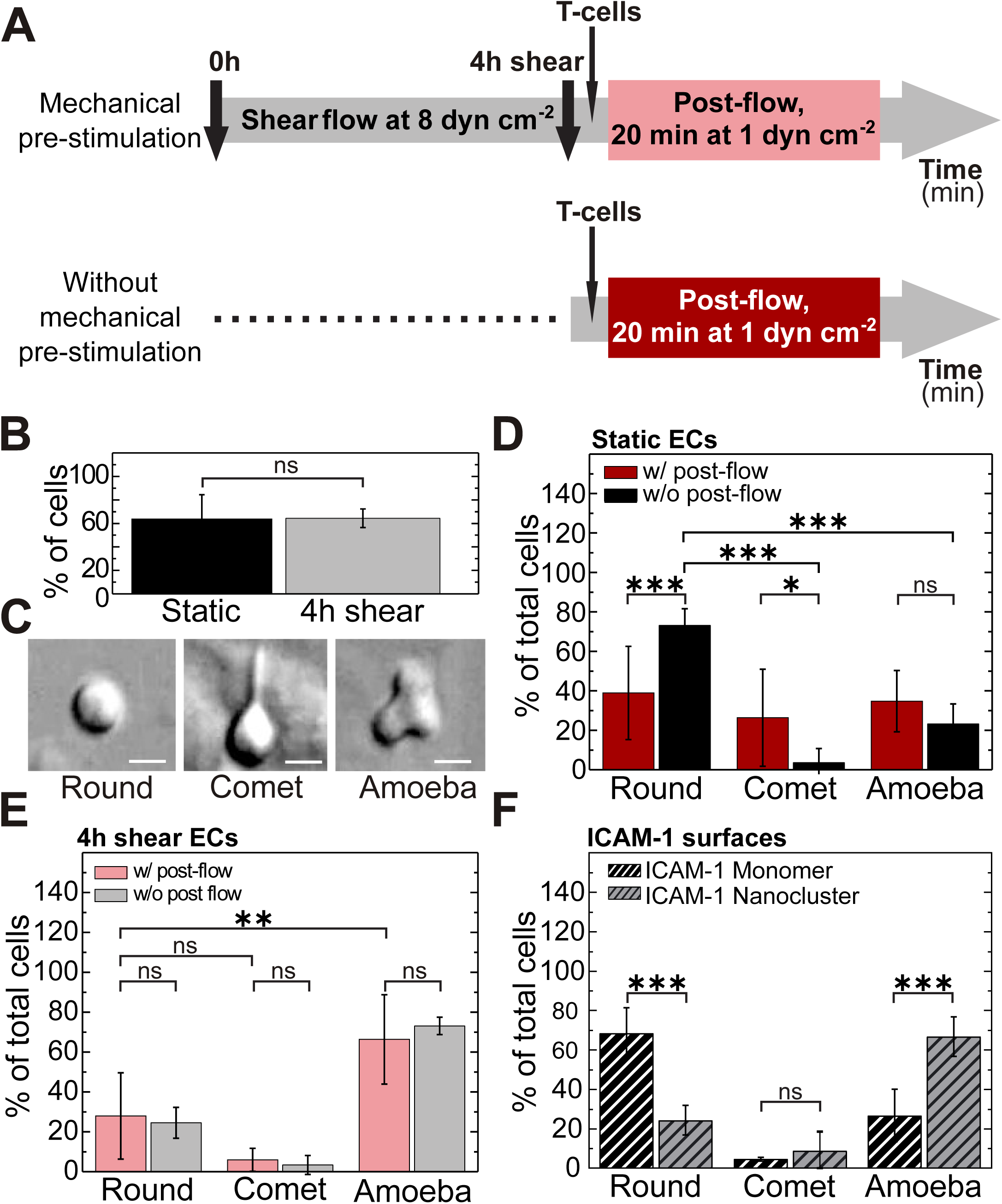
Prolonged shear stimulation of ECs promotes a pro-migratory phenotype on T-cells. (A) Scheme showing the characteristic time scales and course of the experiments with (*top*) and without shear-flow pre-stimulation of ECs (*bottom*). (B) Percentage of T-cells that remained adhered to ECs after 20 minutes of post-flow application. Results are mean ± s.d. (n= 450-600 ECs from 5-10 separate experiments per condition). (C) DIC images showing the three main cell morphologies observed: “ round”, “ comet-like” and “ amoeba-like”. Scale bars, 4 μm. (D) Percentage of T-cells with indicated morphology adhered on static ECs (i.e., without mechanical pre-stimulation), with (w/, red) and without (w/o, black) post-flow for 20 minutes. Results are mean ± s.d. (n=40 T-cells from 4 separate experiments per condition). (E) Percentage of T-cells with indicated morphology adhered to 4 hours, shear-flow pre-conditioned ECs, with (w/, light red) and without (w/o, grey) post-flow for 20 minutes. Results are mean ± s.d. (n=60 T-cells from 5-10 separate experiments per condition). (F) Percentage of T-cells with indicated morphology adhered to monomeric or nanoclustered ICAM-1-coated glass substrates. Results are mean ± s.d. (monomeric ICAM-1 substrates: n= 165 cells from 3 independent experiments; nanoclustered ICAM-1 substrates: n=103 cells from 2 independent experiments). *** *p* < 0.001; ** *p* = 0.01; * *p* = 0.05; ns., not significant.

Once subjected to post-flow, 64% of T-cells firmly adhered to ECs. These results were independent on whether ECs had been initially exposed to prolonged shear-flow or not (Fig. 5B), suggesting that the elevated expression levels of ICAM-1 on TNFα-treated ECs are sufficient to support cell adhesion. Adherent T-cells on ECs underwent noticeable morphological changes in time. We mainly observed three different morphologies, which we classified as “ round”, “ comet-like” and “ amoeba-like”, the latter being consistent with a promigratory phenotype (Fig. 5C). A large majority (ca. 73%) of T-cells adhered to static ECs and in the absence of post-flow remained round (Fig. 5D-*black*), while 20 min of post-flow stimulation on these static ECs showed no preference for a particular type of T-cell morphology (Fig. 5D-*dark red*). In strong contrast, the majority of T-cells (> 60%) on shear-flow pre-stimulated ECs exhibited amoeba-like, promigratory morphology, regardless of whether the T-cells had been additionally subjected to post-flow or not (Fig. 5E). These results indicate that prolonged pre-stimulation of ECs with shear flow promotes a more promigratory response of T-cells.

In the case of shear-flow pre-stimulated ECs, as much as 92% of the T-cells (total of 60) that showed migration, had been initially adhered to EC regions upstream of flow (Fig. S8). Interestingly, these regions do coincide with those in which we had observed ICAM-1 enrichment and nanoclustering. On other EC regions, T-cells did not stably adhere and they were rapidly taken away by the flow. These observations suggest that ICAM-1 nanoclustering and its enrichment to regions upstream of flow might be responsible for the more promigratory phenotype of T-cells. Unfortunately, technical limitations associated with the labeling protocol for STORM and/or STED (i.e., requires cell fixation and Abs labeling) and its incompatibility with our current shear-flow system, prevented us from directly correlating ICAM-1 distribution on ECs with T-cell attachment and/or migration. To overcome these limitations and to emulate the ICAM-1 shear-flow results, we assessed instead whether ICAM-1 distribution (monomer or nanoclustered) on glass-coated surfaces would have an impact on T-cell phenotype. We induced ICAM-1 nanoclusters by means of polyclonal Abs against ICAM-1 (van Zanten et al., 2009), characterized them at the single molecule level using low density concentrations and compared them to the signal from monomeric ICAM-1 to ensure the generation of small nanoclusters (Fig. S9). We then prepared ICAM-1 coated surfaces (monomeric *vs*. nanoclustered) at high density coverage and seeded T-cells on top. T-cells adhered to these two types of substrates showed remarkable differences in their phenotype. Indeed, whereas on monomeric ICAM-1 surfaces most of the T-cells remained round, ICAM-1 nanoclustered surfaces promoted a clear promigratory, amoeba-like morphology (Fig. 5F). These results are qualitative similar to those obtained under prolonged shear-flow pre-conditioning of the ECs (Fig. 5E) and strongly indicate that ICAM-1 nanoclustering is sufficient to induce changes on T-cell morphology and promote their migration.

To further investigate T-cell migration across ECs we applied a tracking algorithm to reconstruct trajectories of individual T-cells (Sosa-Costa et al., 2018) as they migrate in the presence of post-flow (Fig. 6A). An additionally custom-made algorithm detected changes in the velocity and time of interaction of T-cells with ECs (Sosa-Costa et al., 2018) (Fig. 6B). Time-trace curves of cell motion in the direction parallel and perpendicular to the flow were separately analyzed and their respective velocities V_y_ and V_x_ were calculated. The total velocity was determined as V = (V_y_^2^ + V_x_^2^)^1/2^. We defined regions of the trajectory with a positive (or negative) slope as positive (or negative) velocities. The total velocity was considered positive (V+) when the cells were moving *along* the flow direction (V_y_ > 0), and negative (V_-_) when the cells moved *against* the flow direction (V_y_ < 0). Likewise, we defined (t_+_) and (t_-_) as the time periods during which the cell interacted with the ECs, moving *along* or *against* the flow direction, respectively. We then quantified the mean values of V_+_, V_-_, t+ and t_-_ using the cumulative distribution function (CDF) for T-cells migrating over pre-stimulated ECs and compared them to those migrating over static ECs (i.e., without mechanical pre-stimulation) (Fig. S10).

**Fig. 6.**
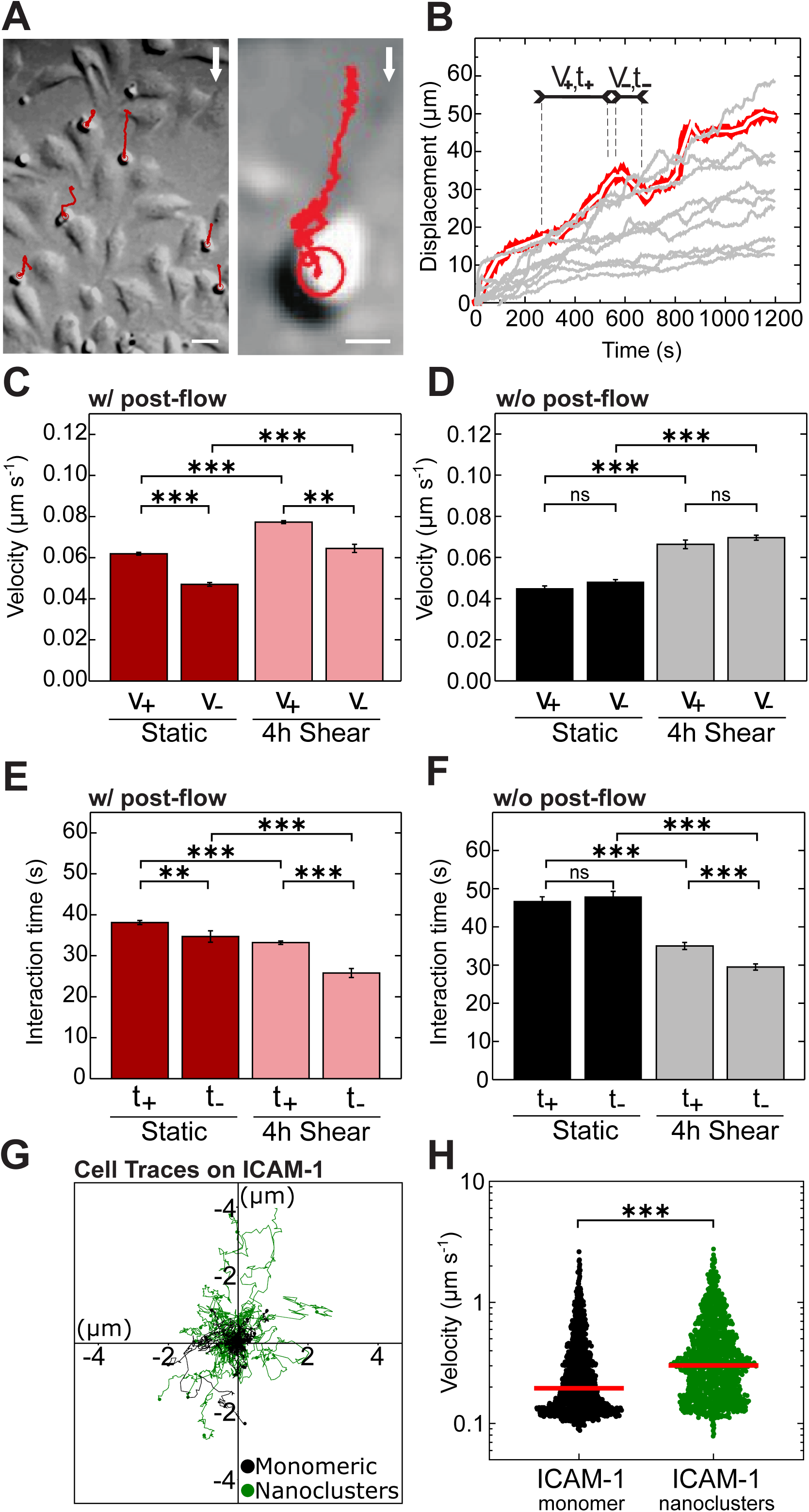
Pre-conditioning of ECs by prolonged shear-flow stimulation influences T-cell migration. (A) Representative DIC (*left*) and zoom-in (*right*) images of T-cells (*small round cells*) migrating in the presence of post-flow over 4 hours shear-flow pre-stimulated ECs (*large elongated cells*). Trajectories of T-cell movement are shown in red. White arrows indicate the flow direction. Scale bars, 20 μm (*left*) and 5 μm (*right*). (B) Characteristic time-trace plots (*grey curves*) of T-cell displacements in the direction parallel to the shear flow. The segmentation algorithm divides the curves into short line segments (*white segments* superimposed over the red line) from which positive and negative slopes, representing the T-cell velocity (*V*_*+*·*−*_) and the time of interaction with ECs (*t*_*+*·*−*_), are determined separately. (C) Velocities of T-cells subjected to post-flow, migrating along (V_+_) or against (V_−_) the flow direction, over static and 4 hours shear-flow stimulated ECs. (D) Similar to (C) but without post-flow. (E,F) Interaction times of T-cells with ECs, with post-flow (E) and without post-flow (F). Results are mean ± s.e.m. (n=60 trajectories per condition from 6-10 independent experiments). (G) Representative single cell trajectories recorded on monomeric (black) or nanoclustered (green) ICAM-1 substrates. (H) Instantaneous velocity of T-cells migrating on the two types of substrates. Red lines correspond to the mean value of the distributions (monomeric surfaces: n= 1989 cell trajectories from 2 independent experiments; nanoclustered surfaces: 1714 cell trajectories from 3 independent experiments). *** *p* < 0.001; ** *p* = 0.01; ns: no significant.

On average, T-cells migrating across shear-flow pre-stimulated ECs moved faster compared to their counterparts moving across static ECs (Fig. 6C). By fitting the experimental data to a single exponential function we estimated an average velocity for cell movement *along* the direction of flow to be V_+_= (77.3 ± 0.7) nm s^-1^ and V_+_= (61.9 ± 0.7) nm s^-1^ for shear-flow pre-stimulated and static ECs respectively (Fig. 6C). These results were independent of whether T-cells were subjected, or not, to post-flow (Fig. 6D) indicating that the increased migration speed was due to the mechanical pre-conditioning of the ECs. Moreover, the mean interaction time between T-cells and ECs exhibited an opposite trend with respect to the velocity, being shorter in the case of shear-flow pre-stimulated ECs, i.e., t_+_= (33.2 ± 0.4)s, as compared to static conditions, i.e., t_+_= (38.1 ± 0.5)s (Fig. 6E) and regardless of post-flow application (Fig. 6F). When T-cells migrated against the flow direction, their velocity was reduced, i.e., V_-_ < V_+_, regardless of whether the ECs were pre-exposed to shear flow or not (Fig. 6C,D). In addition, the interaction time between T-cells and ECs was also shorter for T-cells moving against the flow as compared to those ones moving along the flow (t_-_ < t_+_), independently of the pre-stimulating or post-flow conditions (Fig. 6E,F). Altogether, these results indicate that while the directional motion of T-cells on ECs is driven by the post-flow, their migration behavior strongly depends on the pre-conditioning of ECs by shear flow stimulation: T-cells migrating over mechanically pre-stimulated ECs move at higher speed and make shorter-lived contacts with ECs, as compared to T-cells migrating over static ECs.

Finally, to assess the implications of ICAM-1 nanoclustering induced by prolonged mechanical stimulation on the increased T-cell migration speed, we resourced once more to ICAM-1 coated surfaces. We prepared coated-surfaces with high-density monomeric or nanoclustered ICAM-1 and tracked the mobility of T-cells (final concentration of ∼20μg ml^-1^ of ICAM-1). Consistent with their increased promigratory phenotype on ICAM-1 nanoclustered substrates (Fig. 5F), T-cells migrated significantly faster on these substrates (Fig. 6G,H), once more supporting the notion that ICAM-1 nanoclustering increases T-cell migration. As additional control, we artificially induced ICAM-1 nanoclustering by Ab. cross-linking of living TNFα-stimulated ECs and monitored T-cell migration. Also in these conditions, T-cells migrated significantly faster as compared to control ECs (Fig. S11). Overall, our results strongly indicate that ICAM-1 nanoclustering brought about by shear-flow increases T cell migration.

## DISCUSSION

In this work, we have used confocal and different forms of super-resolution microscopy together with a parallel-plate flow chamber to assess the effect of shear forces on the lateral organization of ICAM-1 on endothelial cells and potential impact on leukocyte migration. Exposure of TNFα-stimulated ECs to shear flow promoted translocation of ICAM-1 to the upstream direction of flow together with the formation of ICAM-1 nanoclusters on the EC membrane *prior* to leukocyte engagement. Moreover, shear-force induced ICAM-1 nanoclustering is actin-dependent. This shear-force induced rearrangement of ICAM-1 and actin directly correlated with altered T-cell migration. Specifically, a more migratory phenotype, faster migration and reduced interaction times of leukocytes with the opposing mechanically pre-stimulated EC surface were observed.

It has been extensively documented that shear-flow induces major re-arrangements of the actin cytoskeleton of ECs with the formation of rich actin fibers along the flow direction (Galbraith et al., 1998). Our super-resolution studies confirmed these changes, and importantly, also showed the formation of actin-patches upstream of flow that localize to newly formed ICAM-1 nanoclusters. Formation of ICAM-1 nanoclusters was actin cytoskeleton dependent and disruption of the actin network led to the dissolution of the nanoclusters. Such local stability of ICAM-1 nanoclusters under shear flow might have been promoted by their anchoring to the actin cytoskeleton by means of different adaptor proteins, such as α-actinin-4 and cortactin (Schaefer et al., 2014). These actin-binding proteins (ABPs) can upregulate directly ICAM-1 clustering and as such, enhance ICAM-1 adhesive function (Schaefer et al., 2014; Schaefer and Hordijk, 2015; Schnoor, 2015). It would be important to further identify which molecular actors are responsible for mechanosensing and the downstream signaling cascade that lead to actin remodeling and ICAM-1 nanocluster formation. Some initial work has been already done along these directions showing that ICAM-1 itself is a force-sensing receptor (Liu et al., 2010), capable of initiating mechanosensitive signaling events, such as recruitment of actin-binding proteins, increase of RhoA activity and ROCK-myosin based contractile forces (Schaefer and Hordijk, 2015). These re-arrangements of the actin machinery could then feedback into ICAM-1, causing its spatial re-distribution on the EC surface. Identifying the pathways involved into the mechanosensing and feedback amplification would provide a great opportunity to target specific steps in the process which would strongly increase our understanding of the molecular mechanisms that underlie the described phenotypes.

Previous studies showed that artificially induced ICAM-1 clustering leads to its immobilization on the EC surface by binding of the ICAM-1 cytoplasmic tail to actin-binding proteins of the ERM and α-actinin families (van Buul et al., 2010). More recently, actin-dependent clustering and ICAM-1 immobilization have been also observed upon dendritic cell maturation (Comrie et al., 2015). Although in our studies we did not assess the mobility of the newly formed ICAM-1 nanoclusters, their strict dependence on the actin cytoskeleton and close proximity to actin patches suggest that these nanoclusters are also immobile. Immobilization of ICAM-1 has major consequences for immune responses that require firm adhesion. Indeed, interruption of ICAM-1 interactions with actinin and members of the ERM family on ECs inhibits the ability of T-cells to undergo diapedesis (Celli et al., 2006; Schnoor, 2015). Moreover, constrained ICAM-1 mobility on dendritic cells promotes T-cell conjugation during the immunological synapse, T-cell homotypic interactions and T-cell proliferation (Comrie et al., 2015). These processes are molecularly regulated by the ability of ICAM-1 to interact with its counter-part integrin receptor LFA-1. Integrin-dependent adhesion is regulated at two levels: affinity (the strength of each individual bond) which depend on the conformational state of the integrin and ability to bind its ligand, and valency (the total number of bonds). The product of affinity and valency provides the avidity and thus strength of the interaction between integrins and their ligands that enable adhesion (Kinashi, 2005). Immobilization and nanoclustering of ICAM-1 will impact on both affinity and valency of LFA-1. Whereas ICAM-1 immobilization will contribute to force-induced conformational changes that lead to LFA-1 activation (Nordenfelt et al., 2016; Schürpf and Springer, 2011), ICAM-1 nanoclustering will increase the number of interactions with LFA-1 molecules, thus augmenting avidity (Bakker et al., 2012; van Kooyk and Figdor, 2000). Our work thus shows that application of shear-forces impact on the spatiotemporal organization of ICAM-1 on ECs, directly affecting the regulatory mechanisms that govern integrin-mediated T-cell adhesion.

Our experiments further showed that T-cell migration depended on prolonged mechanical pre-stimulation of the EC. We observed a more migratory phenotype in T-cells that adhered on mechanically stimulated ECs, and T-cell adhesion was significantly stronger on EC regions upstream of flow, where ICAM-1 levels were found to be the highest. Moreover, T-cells moved faster with shorter interaction periods on ECs pre-exposed to mechanical stress. As ICAM-1 is one of the main ligand receptors implicated in the endothelium-leukocyte interaction by its engagement with LFA-1, we suggest that the changes in the migratory behavior of T-cells should be, at least in part, associated to the spatiotemporal re-organization of ICAM-1 on the EC membrane brought about by shear flow. Indeed, control experiments on monomeric vs. nanoclustered ICAM-1 surfaces conclusively showed that T-cells adhered on these surfaces developed a promigratory profile and migrated faster as compared to monomeric ICAM-1 surfaces. We can qualitatively rationalize our results in the context of a recently proposed motor-clutch model (Bangasser et al., 2017). In essence, such a model predicts that cell velocity at the front edge is regulated by the tight interplay between the strength of engaged integrin-ligand bonds (i.e. clutches) and the stiffness of the substrate. Strengthening of the clutches will slow down actin retrograde flow (Thievessen et al., 2013), thus increasing the effective cell velocity at the front edge. In the context of our experimental results, an increase in ICAM-1 nanoclustering brought about by shear flow will increase the valency to LFA-1 molecules on the T-cell side, reinforcing clutching strength. In addition, as ICAM-1 nanoclusters could become immobilized by its interaction with the actin cytoskeleton (Comrie et al., 2015; van Buul et al., 2010), their engagement with LFA-1 will favor ligand-dependent integrin activation (Nordenfelt et al., 2016; Schürpf and Springer, 2011), further contributing to increase integrin-ligand binding strength. This overall increase will result in a faster advance of T-cells over the ECs, which fully agrees with our experimental observations.

According to the same model, stiffer substrates will induce the earlier break of the integrin-ligand bonds, with a consequent reduction of interaction times (Bangasser et al., 2017). In our experiments, we observed that ICAM-1 nanoclusters formed by shear stress localize to regions enriched with actin. Most probably this limits their spatial diffusion and promotes immobilization (Comrie et al., 2015; van Buul et al., 2010) and effective stiffening on the regions of clutch anchoring. As the effective local substrate becomes stiffer, the bonds between fixed ICAM-1 and the actin cytoskeleton engaged integrins will break faster, reducing the time of interaction between T-cells and ECs as observed in our experiments. The model proposed to connect our overall results is summarized in Fig. 7.

**Fig. 7.**
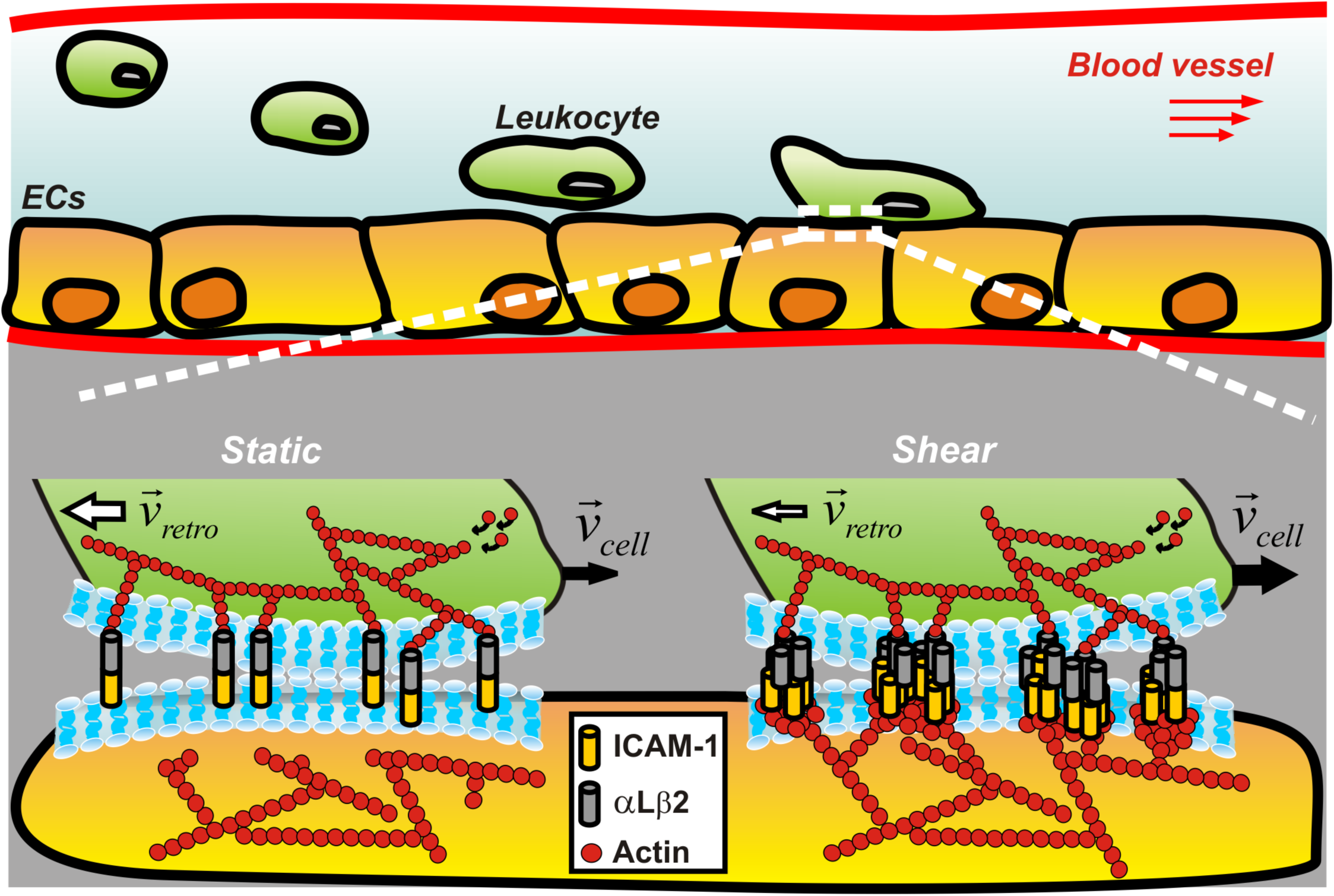
Shear flow-driven actin re-organization promotes ICAM-1 nanoclustering on ECs that affect T-cell migration by strengthening interactions between ICAM-1 and its integrin receptor LFA-1. Under static conditions, leukocyte engagement leads to the formation of bonds between individual ICAM-1 dimers and αLβ2 (LFA-1) counter-integrin receptors. The strength of these bonds regulates leukocyte migration, with the total velocity of the cell edge (*black arrows, V*_*cell*_) being determined by the difference between the speed of the actin cytoskeleton polymerization and the speed of the actin retrograde flow (*white arrows, V*_*retro*_). Shear-flow stimulation of ECs leads to actin-dependent ICAM-1 nanoclustering and most probably, its immobilization on the EC membrane. Both effects enhance the strength of ligand integrin bonds by promoting integrin activation and increasing the avidity of ICAM-1 to LFA-1. This, in turn, reduces the retrograde actin flow rate and increases the total velocity of leukocytes as compared to static conditions. At the same time, as the cell moves forward, the immobilization of ICAM-1 nanoclusters will rapidly lead to ICAM-1-LFA-1 bond breakage, shortening the interaction times between the T-cell and the EC, as compared to static conditions. The size of white and black arrows indicates the resulting velocity, with thicker arrows corresponding to higher speed values.

In summary, our results demonstrate for the first time that shear-forces are sufficient to promote the formation of ligand receptor nanoclusters on the cell surface as a result of force-induced actin-cytoskeleton remodeling. Since many transmembrane receptors interact either directly or indirectly with the actin cytoskeleton, mechanical forces could represent an additional mechanism to orchestrate the lateral organization of a broad range of receptors impacting in their function. In the context of this work, we show that the organization of ligand receptors on the EC membrane influences the migratory behavior of leukocytes. Such a physical mechanism might be of crucial importance for leukocyte-EC interactions during the immune response where the rapid and stable arrest of leukocytes on a vascular bed warrants their precise contact-mediated guidance. In this context, it would be important to assess the role of ICAM-1 polarization and clustering *in-vivo* and its impact in leukocyte transendothelial migration.

## MATERIALS AND METHODS

### Reagents, cytokines and antibodies

RPMI-1640 Dutch modification cell culture medium without phenol red and with glutamine (2 mM) was purchased from Invitrogen. The medium was supplemented with 10% fetal bovine serum (FBS) and 1% antibiotic-antimycotic (AA). Mouse monoclonal primary antibodies against ICAM-1 (anti-CD54) and myosin-II were purchased from BD Pharmingen and Abcam, respectively. Goat-anti-Mouse-AF488 secondary antibody and anti-Rabbit Alexa 647 were from Invitrogen. Purified ICAM-1 Fc Chimera was from R&d systems (Stock concentration 50ug/ml). Poly clonal Anti-ICAM-1 produced in Rabbit was from Santa Cruz (Stock concentration of 200ug/ml). Monoclonal Mouse anti ICAM-1 was from BD bioscience (stock 0.5mg/ml). Human Serum used as blocking medium for the ICAM-1 substrates was from Vitro. For ICAM-1 cross-linking experiments on ECs we used mouse anti ICAM-1 (Clone # BBIG-I1) from R&D systems and secondary antibody from donkey (DK), anti Mouse Alexa 647 from invitrogen. Phalloidin-TRITC, TNFα, CytoD and saponin were from Sigma-Aldrich and phalloidin-AF647 from Life Technologies. Goat-anti-mouse antibody conjugated to Cy3B was prepared in house.

### Cell culture and sample preparation

We used an endothelial cell line, EA.hy926, established elsewhere by fusing primary human umbilical vein cells (HUVECs) with a thioguanine-resistant clone of A549 by exposure to polyethylene glycol (PEG). ECs and Jurkat T lymphoblastoid (a T-cell model) were a gift from Alessandra Cambi (Nijmegen, The Netherlands). In all experiments, ECs were plated on coverslips functionalized with fibronectin (20 μg ml^-1^) and grown to near-confluence in RPMI-1640 medium at 37^°^C humidified atmosphere and in the presence of 5% CO_2_. To mimic inflammatory conditions, samples were activated with TNFα (10 ng ml^-1^) for 20 hours prior to experiments. In experiments with T-cells, cells were collected by centrifugation at 1200 rpm and re-suspended to a concentration of 3 × 10^6^ ml^-1^ in RPMI-1640 medium. In experiments with the actin disrupting agent, a mild concentration of cytoD (5 μg ml^-1^) was flown over 4 hours shear-flow TNFα-stimulated ECs and left in the absence of flow for 20 min. This amount of CytoD and the time of treatment were sufficient to effectively perturb the entire actin cytoskeleton in ECs without causing its total collapse that would lead to cell rounding.

### Microfluidics

Shear flow experiments were performed with a parallel-plate closed flow chamber (FCS2, Bioptechs Inc., Butler, PA). A 40 mm coverslip with a pre-formed endothelial monolayer near confluency was placed on the microaqueduct slide separated by 100 μm-thick gasket with a rectangular surface (1.4 x 2.2 cm). The fully assembled flow chamber was mounted on the stage of an inverted microscope (Olympus IX71) equipped with 20×/0.5 NA air objective and a video camera (CMOS, Thorlabs). A 37“C controlled temperature enclosure surrounding the microscope was used to maintain a constant temperature across the microfluidics. The flow chamber was connected on one end to an automated syringe pump (FCS2 Micro-Perfusion Pump) and on the other end to a medium reservoir. 5% CO_2_-enriched RPMI-1640 cell culture medium of 37°C was perfused at a constant shear flow of 8 dyn cm^-2^ through the chamber. The shear stress of flowing medium *τ*, was calculated using the equation *τ* =(6*Qμ*)/(*wh*^*2*^), where *Q* is the flow rate, *μ* is the viscosity of the fluid, *w* is the width and *h* is the height of gasket.

### T cell adhesion under flow

TNFα-stimulated ECs were exposed to four hours of continuous shear flow at 8 dyn cm^-2^, prior to T-cell adhesion. Next, a suspension of T-cells was flown over ECs using RPMI-1640 perfusion medium at 0.3 dyn cm^-2^. Since non-activated T-cells cannot be captured directly from the flow stream, the flow was stopped to allow T-cells to settle freely on ECs for three minutes. Then, the flow was resumed at 1 dyn cm^-2^ (post-flow). T-cell movement was recorded for 20 minutes by differential interference contrast microscopy (DIC) at 3 frame s^-1^ using a 10×/0.25 NA air objective. In static experiments, T-cells were flown over ECs that were not subjected to shear flow beforehand. Additionally, as control, T-cells were flown over static or shear-flow pre-stimulated ECs, without post-flow application. In these cases, T-cells migrated randomly and over short distances (Fig. S12).

### Preparation and single molecule characterization of monomeric and nanoclustered ICAM-1 substrates

ICAM-1 nanoclusters were prepared following a protocol as reported by van Zanten et al 2009. Briefly, purified ICAM-1-Fc was mixed with polyclonal antibodies in 1% human serum at a final volume of 135 μl (final concentrations of 19 μg ml^-1^ and 20 μg ml^-1^ respectively) and placed in an incubator at 37°C for one hour. ICAM-1 substrates were then prepared in sterile glass bottom petri dishes. High density homogeneous layers of monomeric ICAM-1 were produced by covering the glass bottom petri dish with 120μl of 19 μg ml^-1^ purified ICAM-1 Fc. High density homogeneous layers of nanoclustered ICAM-1 were produced by covering the glass bottom petri dish with 120 μl of the previously prepared aggregated solution. To characterize the degree of ICAM-1 nanoclustering at the single molecule level, we prepared substrates using the same solutions as mentioned above (either monomeric or nanoclustered) but at much lower concentration (0.475μg ml^-1^). Characterization of the samples was performed using the Nikon TIRF set up by imaging with a 647nm laser at 20%, 20ms exposure time and a pixel size of 160 nm. For this, the low density monomeric sample was incubated with primary mouse anti ICAM-1 at 5 μg ml^-1^ for 45 min. The low density nanocluster samples did not require extra primary antibodies as the secondary antibody was already tagged to the primary antibody used to make the nanoclusters. The samples were washed using PBS and secondary antibodies were used. Donkey (DK) anti-mouse Alexa 647 targeted the anti-ICAM-1 on the monomeric samples, and Donkey (DK) anti-rabbit Alexa 647 targeted the polyclonal anti-ICAM-1 for the nanoclusters.

### T-cell adhesion and migration on ICAM-1 substrates

High density ICAM-surfaces were washed with sterile PBS and then covered with cell media. T-cells were added and the samples were placed in the microscope incubation chamber for 10 mins for cells to settle. 20min videos were then taken at 37°C and 5% CO_2_ using a 10x air objective at a frame rate of 3 frames per second. After each 20min video the objective was changed for a 20x objective and cells were imaged for a further 10 mins with the same frame rate. These 10min videos were then used for the phenotype classification experiments.

### Preparation of ECs with antibody-crosslinked ICAM-1 and T-cell adhesion

TNFα-stimulated monolayers of ECs were prepared as indicated above. To artificially generate ICAM-1 clusters on ECs we followed a protocol close to the one described by Millan et al. 2006. In brief, after the stimulation with TNFα, ECs were washed gently with sterile PBS, added 1μg ml^-1^ of polyclonal ICAM-1 antibody diluted in cell media and returned to incubator for 45 minutes at 37°C.ECs were again washed twice with PBS and the secondary antibody to crosslink ICAM-1 (1μg ml^-1^) was added for 20minutes at 37°C. To TNFα-stimulated and ICAM-1 cross-linked ECs monolayers, T cells were then added and allowed to settle for 5-10 minutes. 20min videos were then taken using a 10x air objective at a frame rate of 3 frames per second.

### Immunofluorescence labeling

Resting (without TNFα-stimulation), TNFα-stimulated, static (without mechanical pre-stimulation) and/or shear-flow exposed ECs were washed with warm PBS, fixed in ambient conditions with paraformaldehyde (PFA, 2%) for 15 minutes and immediately stained. Samples were incubated with blocking agents (3% bovine serum albumin, BSA; 10 mM glycine and 1% human serum, HS) for 30 minutes and washed. Next, cells were incubated with primary antibody against ICAM-1 (5 μg ml^-1^) washed and incubated further with the appropriate secondary antibody (2 μg ml^-1^) for 30 minutes. In all cases, cells were stained simultaneously for F-actin with phalloidin-TRITC (0.25 μg ml^-1^) for 40 minutes, after initial permeabilization of the cell membrane with saponin (0.05%) for 10 minutes. Static ECs were double stained for ICAM-1 or myosin-II and F-actin and used as a control. All individual labeling steps were carried out at room temperature. Samples for STORM imaging were immunostained as described above using phalloidin-AF647 for labeling the actin cytoskeleton and antibody conjugated Cy3B for ICAM-1 staining, after initial permeabilization of the cell membrane with triton (0.002%) for 8 min.

### Confocal microscopy

Fluorescence images of ECs were acquired using a laser-scanning confocal microscope (Eclipse TE2000, Nikon). Samples were excited at 488 nm and 561 nm to visualize ICAM-1 or myosin-II and F-actin, respectively. Crosstalk between detection channels was avoided by the use of appropriate excitation and emission bandpass filters. In addition, fluorescence images of ICAM-1 or myosin-II and F-actin were recorded separately by performing a sequential scanning and merged afterward. Three-dimensional image stacks were obtained by scanning through the *z*-direction in steps of 0.1 µm over a range of 20 µm with a piezo-driven 60×/1.49 NA oil immersion objective.

### Stimulated emission depletion (STED) microscopy

Super-resolution images of ICAM-1 clusters were collected with a commercial CW-STED SP-7 microscope (Leica Microsystems) using 100×/1.4 NA oil immersion objective. Samples were excited with an argon-ion laser (488 nm). Images (1024×1024 pixels) were acquired with 12-bit pixel depth, recorded in resonant scan mode at speed of 8 kHz and averaged over 8 frames with line accumulation set at 8. The STED spatial resolution was ∼90 nm as estimated from the size of isolated spots localized at the glass substrate that represent single fluorescence antibodies.

### Dual color stochastic optical reconstruction microscopy (STORM)

Super-resolution images of actin cytoskeleton and ICAM-1 were acquired with a custom-made STORM microscope setup using a standard imaging buffer (1 M Cysteamine MEA, 0.5 mg mL^-1^ glucose, 5% Glucose oxidase and 40µg mL^-1^ catalase in PBS; mixed at a volume ratio of 80:10:10:1). Samples were excited at 647 nm (actin) and 560 nm (ICAM-1) and reactivation of dyes was done with a laser beam at 405 nm. The emitted light was collected with 100×/1.49 NA oil immersion objective, filtered by a quad band filter set (TRF89902-ET-405/488/561/647 Laser Quad Band Set, Chroma) and imaged with an electron-multiplying CCD camera at an exposure time of 20 ms per frame. Images were analyzed using custom-written software (Insight3; provided by Bo Huang, University of California, San Francisco, CA) by fitting the point spread function (PSF) of individual fluorophores with a simple Gaussian curve in every frame to determine the *x* and *y* coordinates. The STORM spatial resolution was ∼25 nm as estimated by calculating the closest resolved distance between two neighboring microtubules (MTs) stained with Alexa 647 in BSC1-African green monkey kidney epithelial cells, in a separate control experiment.

### Image analysis

#### Quantification of ICAM-1 and actin fluorescent intensity from confocal images

The total fluorescence intensity signal of ICAM-1 was calculated as an integrated density signal extracted from individually outlined ECs using the ImageJ software. The *Z*-projected intensity of 80 frames over 8 μm distance, normalized to the cell surface area, is reported. The percentage of the total ICAM-1 fluorescence intensity signal localized either up- or down-stream of flow was estimated from 50% of the desired cell area and divided by the total intensity calculated over the entire cell. Similarly, the up-middle- and downstream percentages of the total ICAM-1 fluorescent intensity signal were estimated from 33% of the corresponding cell area. In total, between 100-180 ECs from two separate experiments were analyzed per condition. The fluorescent intensity of ICAM-1 localized within patch-like actin structures at the apical cell membrane was calculated by manually outlining the desire actin region and integrating the corresponding ICAM-1 density signal over the selected area. In total, 30 ECs from two separate experiments were analyzed per condition.

#### Quantification of ICAM-1 nanoclustering from STED images

Unprocessed, super-resolution STED images were analyzed using custom-written Matlab software. The analysis was based on a 2D Gaussian fit to the intensity profile of manually selected spots from the images. The fitting procedure determines the spot size taking a full-width-at-half-maximum (FWHM) of the Gaussian fit and its intensity (brightness) is calculated as the average of the background-corrected intensity values over all pixels located within the FWHM. To estimate the degree of nanoclustering, the brightness of the spots on the cell membrane were directly compared to that of individual Abs sparsely located on the glass coverslip, outside the cell regions. In total 30 individual cells from two different experiments were analyzed per condition.

#### Quantification of the number of T-cell adhered to ECs after post-flow application

Phase contrast images of 0.05 μm^2^ were taken before and after post-flow application and the number of adherent T-cells in the field of view was counted manually.

#### Generation of T-cell trajectories

The interaction of individual T-cells with ECs were analyzed frame by frame from DIC images using a custom-written MATLAB software. A cross-correlation function was used to estimate changes in the centroid position of individual T-cells over time as described elsewhere (Nelson et al., 2011). In total, 120 cell trajectories (approximately 60 per condition) collected from 6-10 independent experiments were analyzed. T-cells were classified based on cell shape morphology as “ round” (circular morphology), “ comet-like” (circular morphology but with a visible protrusion at the back of the migrating cell) and “ amoeba-like” (spread morphology in various orientations).

#### Analysis of T-cell migration on ECs

A bottom-up type of algorithm for reconstruction and segmentation of noisy time-traces was used to evaluate changes in the velocity and time of interaction of T-cells with ECs (Sosa-Costa et al., 2018). In brief, the algorithm segments the time-traces using a piecewise linear approximation that allows the reconstruction of time-trace data as consecutive straight lines, from which changes of slopes (velocity) and their respective durations can be extracted. The total cell velocity at each time point of an experiment was calculated as the square root of a sum of velocities analyzed in the direction perpendicular (V_x_) and parallel (V_y_) to the flow. The positive (V_+_) and negative (V_-_) signs were assigned to each velocity value to distinguish between periods where T-cells move *along* or *against* the direction of flow, respectively. Accordingly, the positive (t_+_) and negative (t_-_) times of T-cells interactions with ECs were obtained. Velocity and time distributions are represented as a cumulative distribution function (CDF) and fitted with a single exponential function.

#### Analysis of T-cell migration on ICAM-1 substrates and ICAM-1 cross-linked ECs

The videos taken on a 10x air objective were analyzed first in Fiji and then in Python3 in the following way: First, the Fiji edge detection was used for contrast improvement. A TrackMate plugin was used for generating single tracks of the cell movement. Trajectories were then exported to Fiji/Python3 for calculating the instantaneous velocity between consecutive frames. For quantifying the cell phenotype the videos taken on a 20x air objective were used in order to have increased resolution. The phenotypes were counted manually by eye using the Cell Counter plugin in Fiji (ImageJ).

### Statistical analysis

Data are expressed as mean ± standard deviation (s.d.). Where indicated, the standard error of the mean (s.e.m.) is reported. The statistical significance between the means was analyzed using GraphPad Prism 6. The unpaired two-tailed Student *t*-test was used to determine the statistical differences between two individual data sets. The One-way ANOVA test, followed by the Tukey’s multiple comparison test, was used to compare three or more data sets. The Kruskal-Wallis test, followed by Dunn’s multiple comparison test, was used in the case of data sets with non-Gaussian distribution. The *p*-values are indicated.

## Abbreviations used

ICAM-1: Intracellular Adhesion Molecule-1
ECs: endothelial cells
ECs: human umbilical vein endothelial cells
LFA-1: αLβ2 integrin lymphocyte function-associated antigen
Ab: antibody
TNFα: tumor necrosis factor-α
STORM: stochastic optical reconstruction microscopy
STED: stimulated emission depletion microscopy
CytoD: cytochalasin D
ERM proteins: ezrin, radixin, moesin
CDF: cumulative distribution function.

## ACKNOWLEDGMENT

The authors thank T. van Zanten, E. Gutierrez and F. Campelo for useful discussions, J. Andilla from the SLN facility at ICFO for his support with STED imaging and M. Rivas for technical assistance.

## AUTHOR CONTRIBUTION

I.K.P. and M.F.G-P. conceived and designed the experiments. I.K.P., S.K., A.S-C., N.M. and L.L. performed the experiments, microscopy and analyzed the data. A.S-C. and C.M. designed in-silico tools for data analysis. J.S., K.J.E.B., C.M. and M.F.G-P. contributed to data interpretation and suggested experiments. I.K.P. and M.F.G-P. wrote the paper. M.F.G-P. and M.L. supervised the research. All authors revised the manuscript.

## SUPPLEMENTARY INFORMATION

Supplementary information available online at…

## REFERENCES

Abram, C. L. and Lowell, C. A. (2009). The Ins and Outs of Leukocyte Integrin Signaling. Annu Rev Immunol 27, 339–362.

Bakker, G. J., Eich, C., Torreno-Pina, J. A., Diez-Ahedo, R., Perez-Samper, G., van Zanten, T. S., Figdor, C. G., Cambi, A. and Garcia-Parajo, M. F. (2012). Lateral mobility of individual integrin nanoclusters orchestrates the onset for leukocyte adhesion. Proc Natl Acad Sci USA 109, 4869–4874.

Bangasser, B. L., Shamsan, G. A., Chan, C. E., Opoku, K. N., Tüzel, E., Schlichtmann, B. W., Kasim, J. A., Fuller, B. J., McCullough, S. S. and Odde, R&D.J. (2017). Shifting the optimal stiffness for cell migration. Nat Commun 22, 15313.

Butler, P. J., Norwich, G., Weinbaum, S. and Chien, S. (2001). Shear stress induces a time- and position-dependent increase in endothelial cell membrane fluidity. Am J Physiol Cell Physiol 280, C962–C969.

Celli, L., Ryckewaert, J-J., Delachanal, E. and Duperray, A. (2006). Evidence of a Functional Role for Interaction between ICAM-1 and Nonmuscle alpha-Actinins in Leukocyte Diapedesis. J Immunol 177, 4113–4121.

Chen, W., Lou, J. and Zhu, C. (2010). Forcing switch from short-to intermediate-and long-lived states of the αA domain generates LFA1/ICAM1 catch bonds. J Biol Chem 285, 35967–35978.

Comrie, W. A., Li, S., Boyle, S. and Burkhardt, J. K. (2015). The dendritic cell cytoskeleton promotes T cell adhesion and activation by constraining ICAM-1 mobility. J Cell Biol 208, 457–473.

Constantin, G., Majeed, M., Giagulli, C., Piccio, L., Kim, J. Y., Butcher, E. C. and Laudanna, C. (2000). Chemokines trigger immediate beta2 integrin affinity and mobility changes: differential regulation and roles in lymphocyte arrest under flow. Immunity 13, 759–769.

Davies, P. F., Robotewskyj, A. and Griem, M. L. (1994). Quantitative studies of endothelial cell adhesion. Directional remodeling of focal adhesion sites in response to flow forces. J Clin Invest 93, 2031–2038.

Galbraith, C. G., Skalak, R. and Chien, S. (1998). Shear stress induces spatial reorganization of the endothelial cell cytoskeleton. Cell Motil Cytoskel 40, 317–330.

Hogg, N., Patzak, I. and Willenbrock, F. (2011). The insider’s guide to leukocyte integrin signalling and function. Nat Rev Immunol 11, 416–426.

Imhof, B. A. and Aurrand-Lions, M. (2004). Adhesion mechanisms regulating the migration of monocytes. Nat Rev Immunol 4, 432–444.

Kinashi, T. (2005). Intracellular signalling controlling integrin activation in lymphocytes. Nat Rev Immunol 5, 546–559.

Ley, K., Laudanna, C., Cybulsky, M. I. and Nourshargh, S. (2007). Getting to the site of inflammation: the leukocyte adhesion cascade updated. Nat Rev Immunol 7, 678–689.

Li, N., Yang, H., Wang, M., Lü S., Zhang, Y. and Long, M. (2018). Ligand-specific binding forces of LFA-1 and Mac-1 in neutrophil adhesion and crawling. Mol Biol Cell. 29, 408–418.

Liu, Z., Sniadecki, N. J. and Chen, C. S. (2010). Mechanical forces in endothelial cells during firm adhesion and early transmigration of human monocytes. Cell.Mol. Bioeng. 3, 50–59.

Luo, B. H., Carman, C. V. and Springer, T. A. (2007). Structural basis of integrin regulation and signaling. Annu Rev Immunol 25, 619–647.

Makgoba, M. W., Sanders, M. E., Ginther, L. G. E., Dustin, M. L., Springer, T. A., Clark, E. A., Mannoni, P. and Shaw, S. (1988). ICAM-1 a ligand for LFA-1-dependent adhesion of B, T and myeloid cells. Nature 331, 86–88.

Marlin, S. D. and Springer, T. A. (1987). Purified intercellular adhesion molecule-1 (ICAM-1) is a ligand for lymphocyte function-associated antigen 1 (LFA-1). Cell 51, 813–819.

Millan, J., Hewlett, L., Glyn, M., Toomre, D., Clark, P. and Ridley, A.J. (2006). Lymphocyte transcellular migration occurs through recruitment of endothelial ICAM-1 to caveola- and F-actin rich domains. Nat. Cell Biol. 8, 113–127.

Nagel, T., Resnick, N., Atkinson, W. J., Dewey, Jr. C. F. and Gimbrone, Jr. M. A. (1994). Shear stress selectively upregulates intercellular adhesion molecule-1 expression in cultured human vascular endothelial cells. J Clin Invest 94, 885–891.

Nakajima, H. and Mochizuki, N. (2017). Flow pattern-dependent endothelial cell responses through transcriptional regulation. Cell Cycle 16, 1893–1901.

Nelson, P. C., Zurla, C., Brogioli, D., Beausang, J. F., Finzi, L. and Dunlap, D. (2011). Tethered particle motion as a diagnostic of DNA tether length. Int J Nanomedicine 6, 179–195.

Nordenfelt, P., Elliott, H. L. and Springer, T. A. (2016). Coordinated integrin activation by actin-dependent force during T-cell migration. Nat Commun 7, 13119.

Nourshargh, S. and Alon, R. (2014). Leukocyte migration into inflamed tissues. Immunity 41, 694–707.

Schaefer, A., Riet, J. T., Ritz, K., Hoogenboezem, M., Anthony, E. C., Mul, F. P. J., de Vries, C. J., Daemen, M. J., Figdor, C. G., van Buul, J. D., et al. (2014). Actin-binding proteins differentially regulate endothelial cell stiffness, ICAM-1 function and neutrophil transmigration. J Cell Sci 127, 4470–4482.

Schaefer, A. and Hordijk, P. L. (2015). Cell-stiffness-induced mechanosignaling - a key driver of leukocyte transendothelial migration. J Cell Sci. 128, 2221–2230.

Schnoor, M. (2015). Endothelial actin-binding proteins and actin dynamics in leukocyte transendothelial migration. J Immunol. 194, 3535–3541.

Schürpf, T. and Springer, T. A. (2011). Regulation of integrin affinity on cell surfaces. EMBO J 30, 4712–4727.

Shamri, R. l., Grabovsky, V., Gauguet, J-M., Feigelson, S., Manevich, E., Kolanus, W., Robinson, M. K., Staunton, D. E., von Andrian, U. H. and Alon, R. (2005). Lymphocyte arrest requires instantaneous induction of an extended LFA-1 conformation mediated by endothelium-bound chemokines. Nat Immunol 6, 497–506.

Sosa-Costa, A., Piechocka, I. K., Gardini, L., Pavone, F. S., Capitanio, M., Garcia-Parajo, M. F. and Manzo, C. (2018). PLANT: A Method for Detecting Changes of Slope in Noisy Trajectories. Biophys J 114, 2044–2051.

Staunton, D. E., Dustin, M. L., Erickson, H. P. and Springer, T. A. (1990). The arrangement of the immunoglobulin-like domains of ICAM-1 and the binding sites for LFA-1 and rhinovirus. Cell 61: 243–254.

Thievessen, I., Thompson, P. M., Berlemont, S., Plevock, K. M., Plotnikov, S. V., Zemljic-Harpf, A., Ross, R. S., Davidson, M. W., Danuser, G., Campbell, S. L., et al. (2013). Vinculin-actin interaction couples actin retrograde flow to focal adhesions, but is dispensable for focal adhesion growth. J Cell Biol 202.

Tsuboi, H., Ando, J., Korenaga, R., Takada, Y. and Kamiya, A. (1995). Flow stimulates ICAM-1 expression time and shear stress dependently in cultured human endothelial cells. Biochem Biophys Res Commun 206, 988–996.

van Buul, J. D., van Rijssel, J., van Alphen, F. P. J., Hoogenboezem, M., Tol, S., Hoeben, K. A., van Marle, J., Mul, E. P. J. and Hordijk, P. L. (2010). Inside-out regulation of ICAM-1 dynamics in TNF-alpha-activated endothelium. PLOS ONE 5, e11336.

van Kooyk, Y. and Figdor, G. (2000). Avidity regulation of integrins: the driving force in leukocyte adhesion. Curr Opin Cell Biol 12, 542–547.

van Zanten, T. S., Cambi, A., Koopman, M., Joosten, B., Figodr, C.G. and Garcia-Parajo, M.F. (2009). Hotspots of GPI-anchored proteins and integrin nanoclusters function as nucleation sites for cell adhesion. Proc Natl Acad Sci USA 106, 18557–18562.

van Zanten, T. S., Gómez, J., Manzo, C., Cambi, A., Buceta, J., Reigada, R. and Garcia-Parajo, M. F. (2010). Direct mapping of nanoscale compositional connectivity on intact cell membranes. Proc Natl Acad Sci USA 107, 15437–15442.

Wojciak-Stothard, B. and Ridley, A. J. (2003). Shear stress-induced endothelial cell polarization is mediated by Rho and Rac but not Cdc42 or PI 3-kinases. J Cell Biol 161, 429–439.

Yamamoto, K. and Ando, J. (2013). Endothelial cell and model membranes respond to shear stress by rapidly decreasing the order of their lipid phases. J Cell Sci 126, 1227–1234.

Yang, Y., Jun, C. D., Liu, J. H., Zhang, R., Joachimiak, A., Springer, T. A. and Wang, J. H. (2004). Structural basis for dimerization of ICAM-1 on the cell surface. Mol Cell 14, 269–276.

